# Coral long-term recovery after bleaching: implications for sexual reproduction and physiology

**DOI:** 10.1101/2024.04.09.588789

**Authors:** JL Padilla-Gamiño, E Timmins-Schiffman, EA Lenz, SJ White, J Axworthy, A Potter, J Lopez, F Wang

## Abstract

This study examined the long-term impacts of coral bleaching on the reproduction and physiology of *Montipora capitata*, a dominant reef-building coral in Hawaiʻi. We monitored bleached and non-bleached colonies during and after a natural coral bleaching event in 2014 and analyzed reproductive traits and transcriptomic signatures eight months later. Our study shows that non-bleached and bleached colonies successfully produced gametes. Colonies that bleached had smaller oocytes, and development was slower than in colonies that did not bleach. Corals with different vulnerabilities to bleaching exhibited distinct transcriptomic responses eight months after a bleaching event. Those more prone to bleaching showed suppression of transcripts associated with sperm motility, calcification, and immunity. We found distinct transcriptomic signatures between fringing and patch reefs, suggesting local adaptation and/or acclimatization. To conserve coral reefs and better understand how they will be affected by future heat stress, we need to track which colonies survive and examine how their physiological and reproductive processes are impacted in the short- and long-term. This is critical as consecutive bleaching events become more frequent, and corals have less time to recover. Our study provides valuable molecular and reproductive data that can be used for conservation and management purposes. This information can help us identify signs of coral vulnerability and resilience to bleaching, project how future bleaching events will affect coral reproduction, determine which traits are most at risk, and assess which sites are more likely to be compromised.

## INTRODUCTION

Ocean warming is the leading cause of coral mortality due to bleaching events worldwide (Hughes et al., 2017). During bleaching events, corals lose their symbiotic relationship with dinoflagellate endosymbionts (Family Symbiodiniaceae, LaJeunesse et al., 2018), which provide them with color and energy from photosynthesis (Muscatine et al., 1984). Without this energy, coral health is compromised with significant consequences for survival, growth, and reproduction (Coles & Brown, 2003; Hoegh-Guldberg, 1999). In the last decade, coral bleaching events have increased in frequency, duration, and severity (van Hooidonk et al., 2013, Lough and Hughes 2018), causing ecological, socioeconomic, and cultural losses (Chaijaroen, 2022; Pratchett et al., 2008).

Coral response to thermal stress varies depending on temperature tolerance, environmental factors (Hoegh-Guldberg et al., 2007; Hughes et al., 2017), genetics of the host and symbiont, epigenetics, and microbial community composition (Baird et al., 2009; Edmunds, 1994; Fitt & Warner, 1995; Grottoli et al., 2014; Marshall & Baird, 2000; Rosenberg et al., 2009; van Oppen et al., 2015). During and after thermal stress, corals either die or acclimatize and recover. Acclimatization and recovery occur via changes in physiology (e.g., plasticity) manifested in shifts in gene and protein expression, epigenetic changes, symbiont shuffling changes in dominant symbiont after bleaching (Baker, 2001; Hoegh-Guldberg, 1999), and heterotrophic plasticity (Grottoli et al., 2006; Rodrigues et al., 2008). Corals sometimes divert energy from reproductive processes and allocate it towards recovery (Michalek-Wagner & Willis 2001, Rodrigues & Padilla-Gamiño, 2022, Jeffe et al. 2023).

Acclimatization mechanisms associated with thermal stress response and recovery in the coral host may occur at different temporal scales (van Woesik et al., 2022). Gene expression changes can occur within minutes (Thomas et al., 2019; Traylor-Knowles, Rose, & Palumbi, 2017; Traylor-Knowles, Rose, Sheets, et al., 2017), whereas symbiont reacquisition, reallocation of energy, and reproductive recovery can occur over months or years (Rodrigues et al., 2008; van Woesik et al., 2022). Coral thermal stress response can involve hundreds of transcripts involved in apoptosis, oxidative stress, transcription regulation and redox regulation (Bellantuono et al., 2012; Kenkel et al., 2013; Maor-Landaw & Levy, 2016; Thomas et al., 2019; Williams et al., 2021; Zhang et al., 2022). Examining transcriptomic signatures before, during, and after stress can help us to understand coral resilience, identify biomarkers of thermal tolerance, assess physiological health, and better understand the ability and mechanisms of individuals to respond to and recover from environmental stress (Pinzón et al., 2015; Seneca & Palumbi, 2015; Thomas et al., 2019). To date, most coral transcriptomic studies have focused on short-term responses during and after bleaching and very few studies have focused on gene expression signatures associated with long-term recovery in the field (McLachlan et al., 2020; Pinzón et al., 2015; Thomas et al., 2019; Thomas & Palumbi, 2017).

Little is known about how corals can continue long-term and expensive energy processes such as reproduction after stress (Cox, 2007; Rodrigues & Padilla-Gamiño, 2022). This information is critical to project both the short-term reproductive potential of survivors and the longer-term population persistence in the future with more severe and frequent bleaching events. Observations of coral reproduction following natural bleaching events reveal a varied response among and within coral species, suggesting different physiological mechanisms during recovery. Consequences of reproduction due to bleaching stress include a decrease in gametogenesis, oocyte production, oocyte size, sperm motility, spermary abundance, fecundity, and spawning (Baird & Marshall, 2002; Cox, 2007; Hagedorn et al., 2016; Mendes & Woodley, 2002; Omori, 2011; Szmant & Gassman, 1990). There are species that either abort (Jokiel et al., 1985) or substantially decrease oocyte and/or planula (i.e., free-swimming larva) production following exposure to warm seawater temperatures (Airi et al., 2014; Ward & Harrison, 2000). *Orbicella annularis* colonies that had visibly recovered from bleaching (in 3-6 months) followed a normal reproductive cycle, while colonies that remained visibly bleached for more than 8 months did not (Szmant & Gassman, 1990). A two-year reduction in gonad development was observed in bleached corals of *O. annularis* (Mendes & Woodley, 2002) and an eleven-year study (2002-2013) by (Levitan et al., 2014) on the *Orbicella* species complex found that a severe coral bleaching event reduced the likelihood of spawning in both visibly bleached and non-bleached corals for up to four years. In contrast, other species have the capacity to continue sexual reproduction despite bleaching (Armoza-Zvuloni et al., 2011; Cox, 2007; Padilla-Gamiño & Gates, 2012; Szmant & Gassman, 1990).

To better understand long-term coral physiological recovery after a bleaching event, we examined transcriptomic signatures and reproductive traits in colonies of *Montipora capitata* with different thermal tolerance (bleaching susceptibility). *M. capitata* is an important reef builder in Hawai’i and a dominant species in Kāneʻohe Bay, Oʻahu (Edmondson 1929, Maragos 1972, Jokiel 1991). In 2014, Hawaiian reefs experienced an extensive bleaching event with ∼45% of corals affected by bleaching, and in some areas of Kāneʻohe Bay, bleaching reached up to 80-100% (Bahr et al., 2015, 2016; Ritson-Williams & Gates, 2020). From September to October 2014, the sea surface waters in Kāneʻohe Bay exceeded 28 °C, which is the critical threshold temperature for coral bleaching in Hawaiʻi (Bahr et al., 2015; Jokiel & Brown, 2004), and in this period, there were 6 days where the mean temperature was 30 °C and above in Kāneʻohe Bay (Ritson-Williams & Gates, 2020). During the bleaching event, some colonies bleached entirely in small reef areas, while others kept their coloration and seemed unaffected (Bahr et al., 2015).

To assess the impacts of bleaching on the fitness of *M. capitata,* we examined reproduction and gene expression differences in bleached and non-bleached colonies eight months after the 2014 bleaching event (when gametogenesis was completed). We obtained colonies from fringing and patch reefs to test whether (1) transcriptomic signatures differ between corals that bleached and did not bleach, (2) fecundity, oocyte size, and gametogenesis are affected by bleaching, and (3) environment influences coral transcriptomic signatures and reproductive performance.

## METHODS

### Field Sites and Sample Collections

During the 2014 bleaching event in Kāneʻohe Bay (Oʻahu, Hawaiʻi, Fig. 1), we tagged bleached (completely whitened) and non-bleached coral colonies of *Montipora capitata.* This species is a simultaneous hermaphrodite that releases egg-sperm bundles in the summer months close to the new moon (Heyward 1986). Colonies ranging in size from 0.5 - 2 m in diameter were tagged on the leeward side of two patch reefs in October 2014 (Reef 44: 21°28.595’ N, 157°50.013’ W and HIMB: 21°26.159 N, 157°47.489’W) and on two fringe reefs in December 2014 (K4: 21°26 684 N, 157°48 362 W and K5: 21°28 032 N, 157°49 977 W).

**Figure 1.**
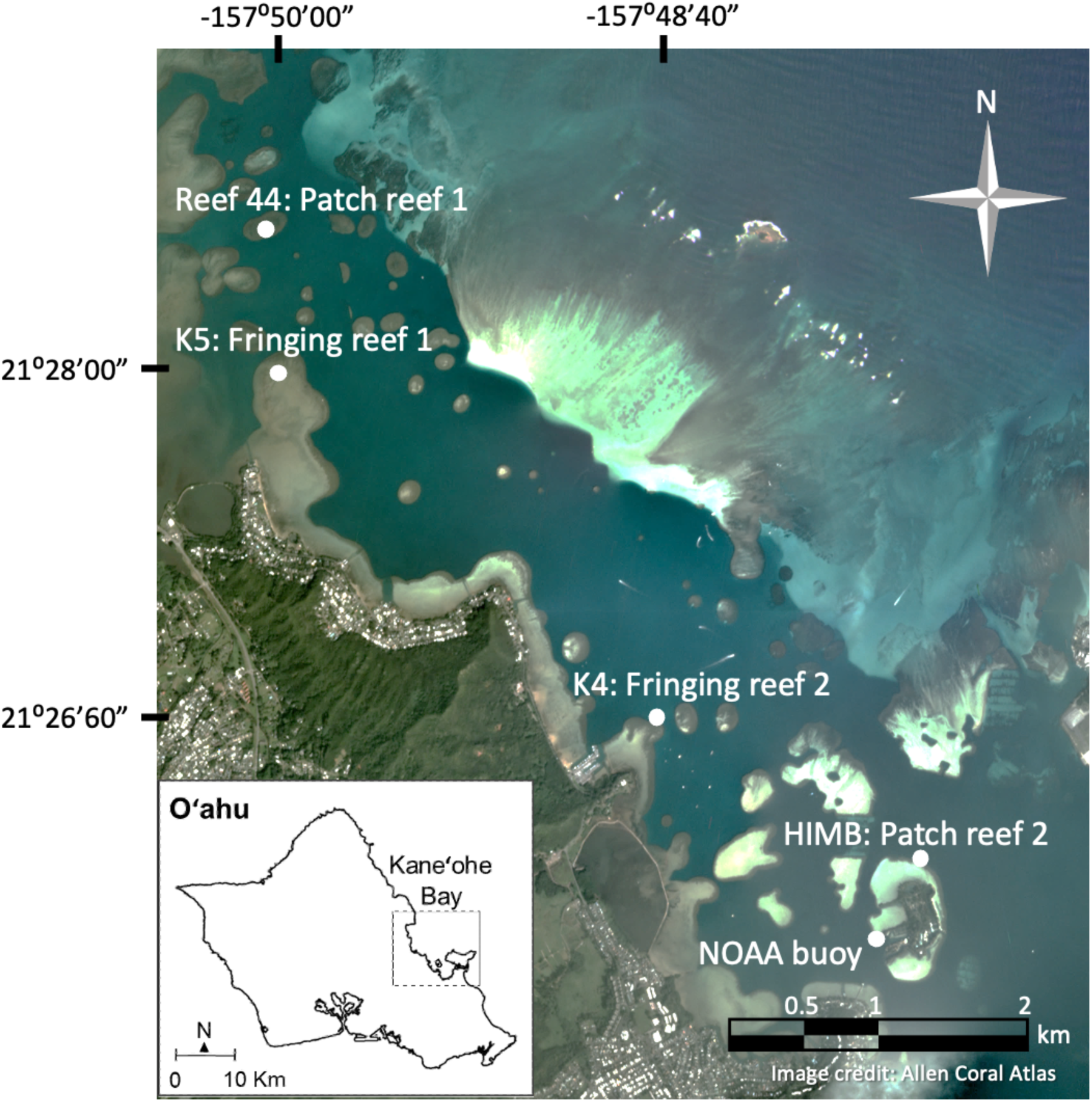
Map of study sites (Reef 44, K4, K5, and HIMB) at Kāneʻohe Bay, Hawaiʻi.

On June 13-14th, 2015, we collected 2 - 4 cm fragments from the tagged colonies (n = 6-10) to assess physiology (gene expression) and reproduction (gamete development and fecundity) in corals that bleached and did not bleach during the thermal stress event of 2014. Collections occurred 2 to 3 days before the new moon when spawning is expected for *M. capitata*, and eight months after the thermal stress event (Fig. 2).

**Figure 2.**
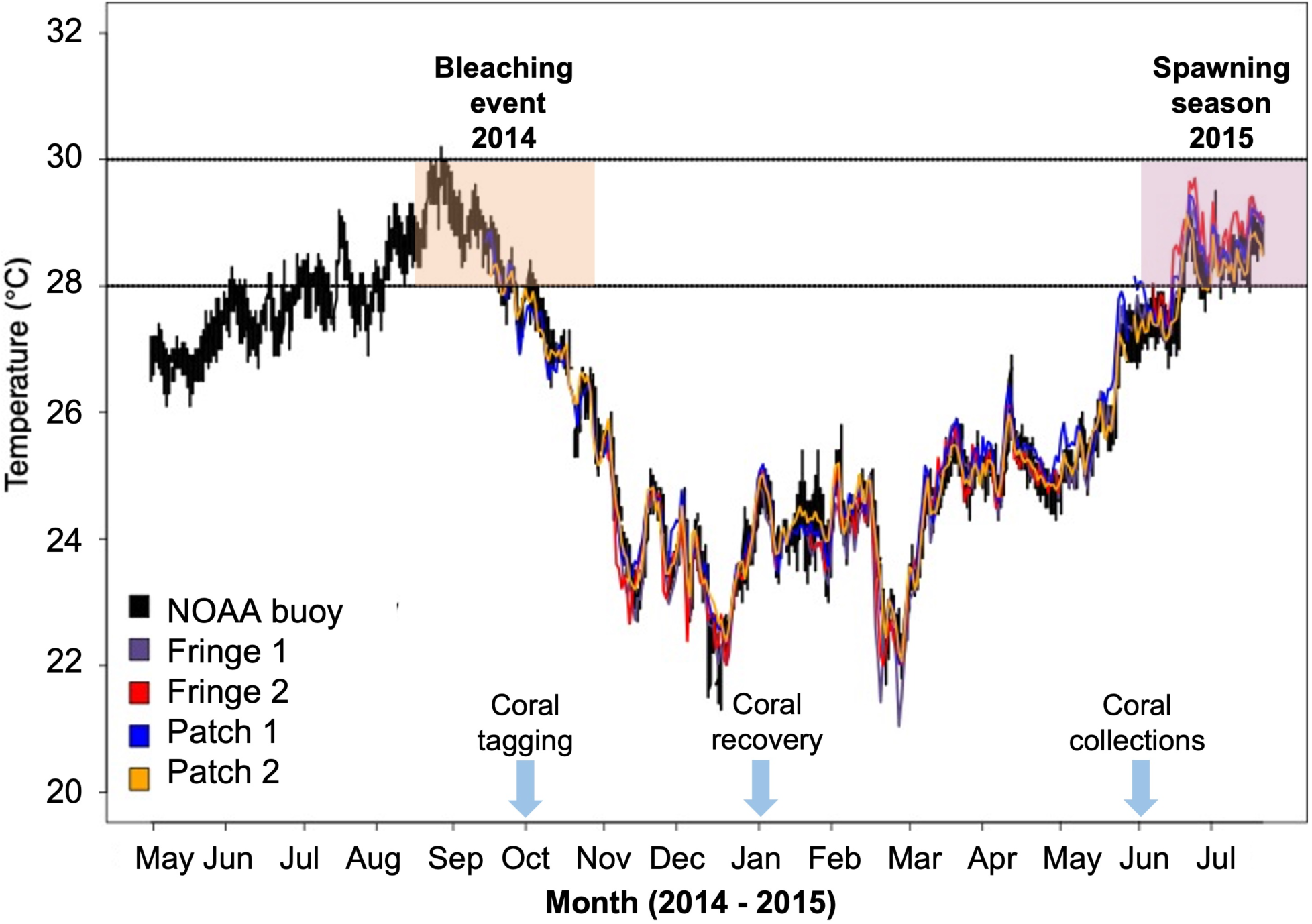
Thermal history in Kāneʻohe Bay at two fringing reefs (K4 and K5), two patch reefs (HIMB and reef 44) and NOAA buoy station located near Moku o Loe Island (HIMB). Figure shows the timing when we tagged vulnerable and susceptible colonies (bleached or non-bleached), when colonies recovered and when we collected the samples to assess reproductive characteristics.

Corals in the bay’s northern part were collected at fringing reef K5 and patch reef 44. Fringing reef K5 was located near the He‘eia fishpond, and the output of the Waihe‘e and Kahalu‘u streams and reef 44 was located near the Shipping (NW) channel. In the southern part of the bay, corals were collected at fringing reef K4 and patch reef HIMB close to the Sampan (SE) channel (Fig. 1).

Temperature data was recorded using HOBO loggers (Onset Computer; accuracy ± 0.2°C) recording every 15 minutes. Temperature from reef 44 and HIMB (patch reefs) was collected from October 2014 to August 2015 and temperature from K5 and K4 stations was collected from November 2014 to August 2015 (Fig. 1, Table 1). Light data was recorded every 15 min from June to October 2015 using Odyssey Photosynthetic Active Radiation (PAR) sensors. Minimum water temperatures occurred in January and March throughout the bay (Fig. 2). Maximum water temperatures occurred in July-August and were higher in fringing reefs compared to patch reefs (∼29°C and 28°C respectively, Table 1). Overall, temperature fluctuates more in the fringing reefs than the patch reefs (Table 1). K5 (fringing reef in the northern site) had the highest variability in temperature and light conditions (Table 1), and the lowest light levels of all the sites. K5 is found near the He’eia fishpond and the output of the Waihe’e and Kahalu’u streams and has higher sedimentation rates than K4 throughout the year (Padilla-Gamiño et al., 2014).

### Histology

Samples were fixed in the field immediately after collection using buffered zinc formalin fixative (Z-fix, Anatech Ltd) diluted in 0.2 - µm filtered seawater (1:4). After 48-72 hours, corals were transferred into 70% ethanol (diluted in DI water) until processing. Fixed coral samples were decalcified using a buffered 10% hydrochloric acid solution for 24 h. Tissue samples were processed for histological examination according to Padilla-Gamino et al. (2014). Developmental stages of oocytes and spermocytes were identified based on cell size and morphology criteria (Szmant-Froelich *et al.,* 1980; Szmant-Froelich, 1985, Padilla-Gamiño et al., 2014). Feret’s Statistical Diameter was used to estimate the size of the oocytes using ImageJ Software V. 1.42 (Abramoff et al., 2004). Size measurements were performed only on oocytes sectioned through the nucleus in order to standardize measurements to the widest axis of oocytes to ensure no duplicate measurements (Davis, 1982; Parker et al., 1997). Oocyte size and spermatocyst development were compared with previously reported data from the same species and location in a year when no coral bleaching was reported (Padilla-Gamiño et al., 2014). We compared results of our histological analysis with those from colonies from the same region, collected and fixed on June 4, 2009 (18 days before the new moon) (Padilla-Gamiño et al., 2014).

### RNA extractions

Samples were immediately frozen after collection using liquid nitrogen and then stored in a - 80°C freezer until processing. Ambion RNAqueous Total RNA Isolation Kit (AM1912) was used to extract RNA. The coral sample was crushed in 1 mL of lysis buffer using a razor blade and further homogenized by passing it through a 25 g syringe needle several times. The samples were then centrifuged for three minutes at top speed to remove debris, and the supernatant was transferred to a new 1.5 mL tube. Following the protocol of the RNAqueous Kit, RNA in the supernatant was isolated and eluted with 50 μL preheated Elution Solution. The quantity and quality of coral RNA samples were evaluated by an Agilent Bioanalyzer.

### Library preparation and sequencing

Library preparation and sequencing were performed at the Institute for Integrative Genome Biology at the University of California Riverside. Stranded libraries were prepared with 100 ng of total RNA with the TruSeq Stranded mRNA Library Prep for NeoPrep Kit (Illumina, Inc.) per the manufacturer’s instructions. Indexed libraries were pooled and run on a HiSeq4000 to generate 100bp paired-end reads for each library. All FastQ files are available in the NCBI Short Read Archive (NCBI BioProject: PRJNA735085; SRR14729868 - SRR14729894).

### Transcriptome assembly and annotation

FastQ file quality trimming, transcriptome assembly, quality assessment, and annotation are described in the supporting information.

### Statistical analysis

#### Reproductive status and bleaching

The impacts of previous bleaching condition and reef type on reproductive traits were investigated using generalized linear models with the lme4 (v1.1-27.1) package for generalized linear mixed-effects models and the nlme (v 3.1-155) package for linear mixed-effects models in R following the methods of (Leinbach et al., 2021). Effects were considered significant with p-value < 0.05. To compare oocyte and spermatocyst stages between a non-bleaching year (2009) and a bleaching year (2015), gamete stages for the two years were combined for the fringe reef only and analyzed as described above.

#### Transcriptomics

Outlier samples and genes were screened out of the dataset using the goodSamplesGenes command in the WGCNA package in R (Langfelder & Horvath 2008). Nonmetric multidimensional scaling (NMDS) ordination was performed on the log(x+1)-transformed reads from the reduced transcriptome using a Bray-Curtis dissimilarity matrix in the vegan package (Oksanen et al. 2017) in R. Analysis of similarity (ANOSIM) was performed on the row-standardized data in a Bray-Curtis dissimilarity matrix to compare significant transcriptomic differences between groups based on the comparisons of: 1) all factors (bleaching and reef type); 2) reef type only; and 3) bleaching only. Discriminant analysis of principal components (DAPC) was used to reduce intra-group variability and find the underlying transcript clusters that differentiated groups based on bleaching and site of origin. Based on xval scores, 2 PCs were maintained for the DAPC analysis and clusters with a loading value of at least 0.01 were considered significant.

Weighted gene correlation network analysis (WGCNA) was performed on the clustered transcripts from the cd-hit pipeline following the instructions for working with a large dataset in R using a blockwise analysis. WGCNA modules were considered significantly correlated with reef type or bleaching status if the absolute value of the correlation coefficient was at least 0.7.

The counts matrix from corset was used as the RNASeq matrix for differential expression analysis in edgeR. DGE comparisons were performed on 1) bleached and non-bleached corals from the fringing reef; 2) bleached and non-bleached corals from the patch reef; 3) bleached and non-bleached across reef types; and 4) patch and fringing, regardless of bleaching status. Transcripts were considered differentially expressed with FDR < 0.05. Heatmaps of the differentially expressed genes were made in pheatmap in R.

Gene Ontology (GO) enrichment analysis was performed on groups of differentially expressed genes using compGO <u>(Timmins-Schiffman et al. 2017)</u> (project-specific enrichment portal: https://meta.yeastrc.org/compgo_emma_montipora_large/pages/goAnalysisForm.jsp).

## RESULTS

### Reproduction (gamete presence, fecundity, gametogenesis)

The probability of an *M. capitata* colony containing gametes varied significantly with reef type; colonies found at fringing reefs were more likely to have gametes present than those at patch reefs (p=0.014, Fig. 3). Bleaching history had no impact on gamete presence. At fringing reefs, all colonies that bleached had gametes, and only a few colonies that did not bleach had no gametes.

**Figure 3.**
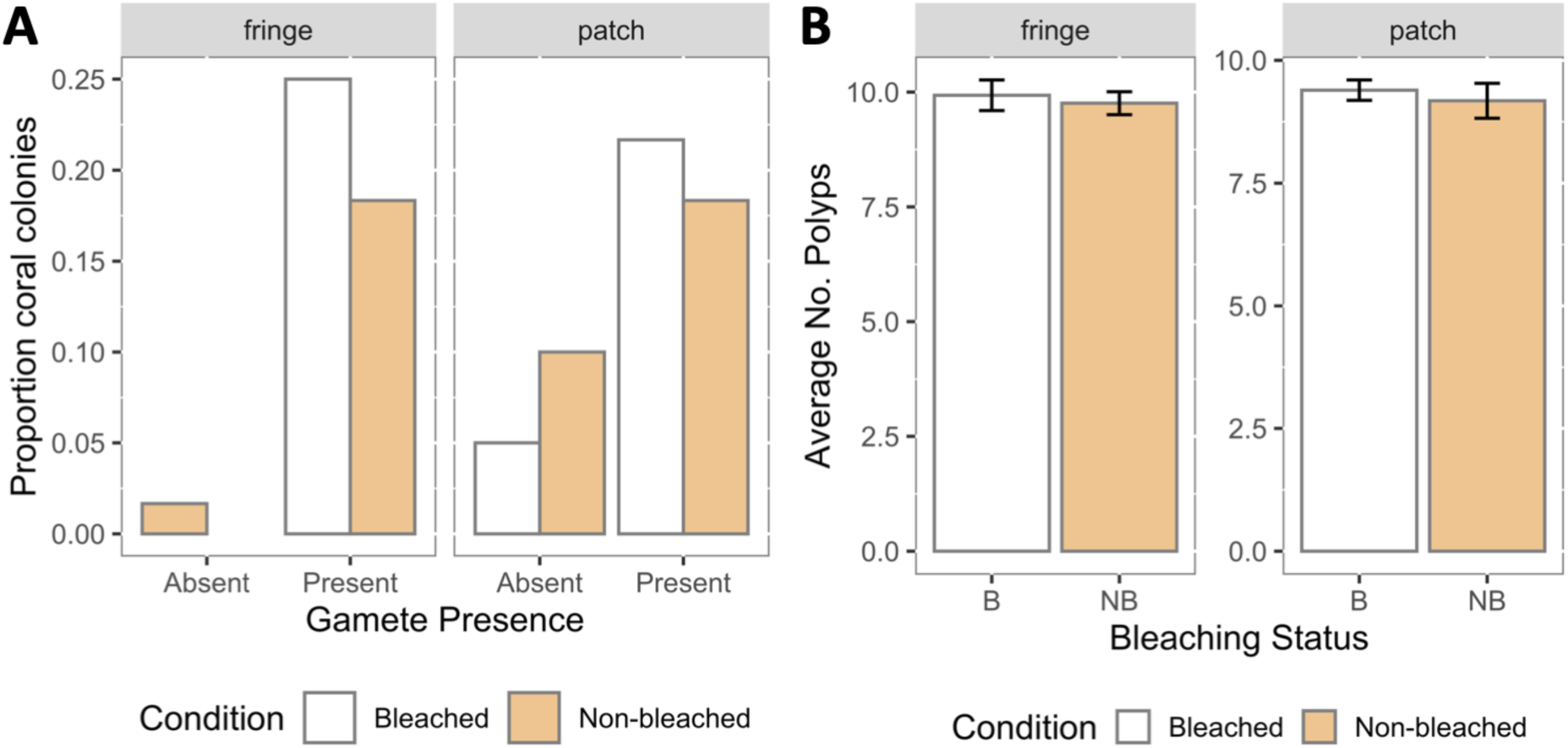
Gamete presence and fecundity in bleached and non-bleached colonies of *Montipora capitata*

There was no difference in fecundity (number of polyps) between previously bleached and non-bleached corals; neither was there a difference in fecundity based on reef type (Fig. 3).

There was a significant difference in oocyte size (Feret diameter) based on bleaching history (p=0.0005) and reef type (p=0.024). Corals that did not bleach had larger oocyte size (Fig. 4) and slightly larger oocytes were found in fringing reefs than in patch reefs (∼1.5% difference). Previously bleached corals from fringing reefs had an average Feret diameter of 272.56 µm, compared to 294.45 µm in non-bleached corals (∼7% difference). In corals from the patch reefs, bleached corals had an average Feret diameter of 279.41 µm, while non-bleached corals had an average Feret diameter of 296.17 µm (∼6% difference). Oocytes of bleached and non-bleached colonies in our study were smaller (19% and 13%, respectively) compared to oocytes from a period (2008-2009) when no bleaching was reported (Padilla-Gamiño et al., 2014). Bleaching status (p=0.000), and year (p=0.0163) were significant predictors of oocyte size when analyzing the 2015 and 2009 data together.

**Figure 4.**
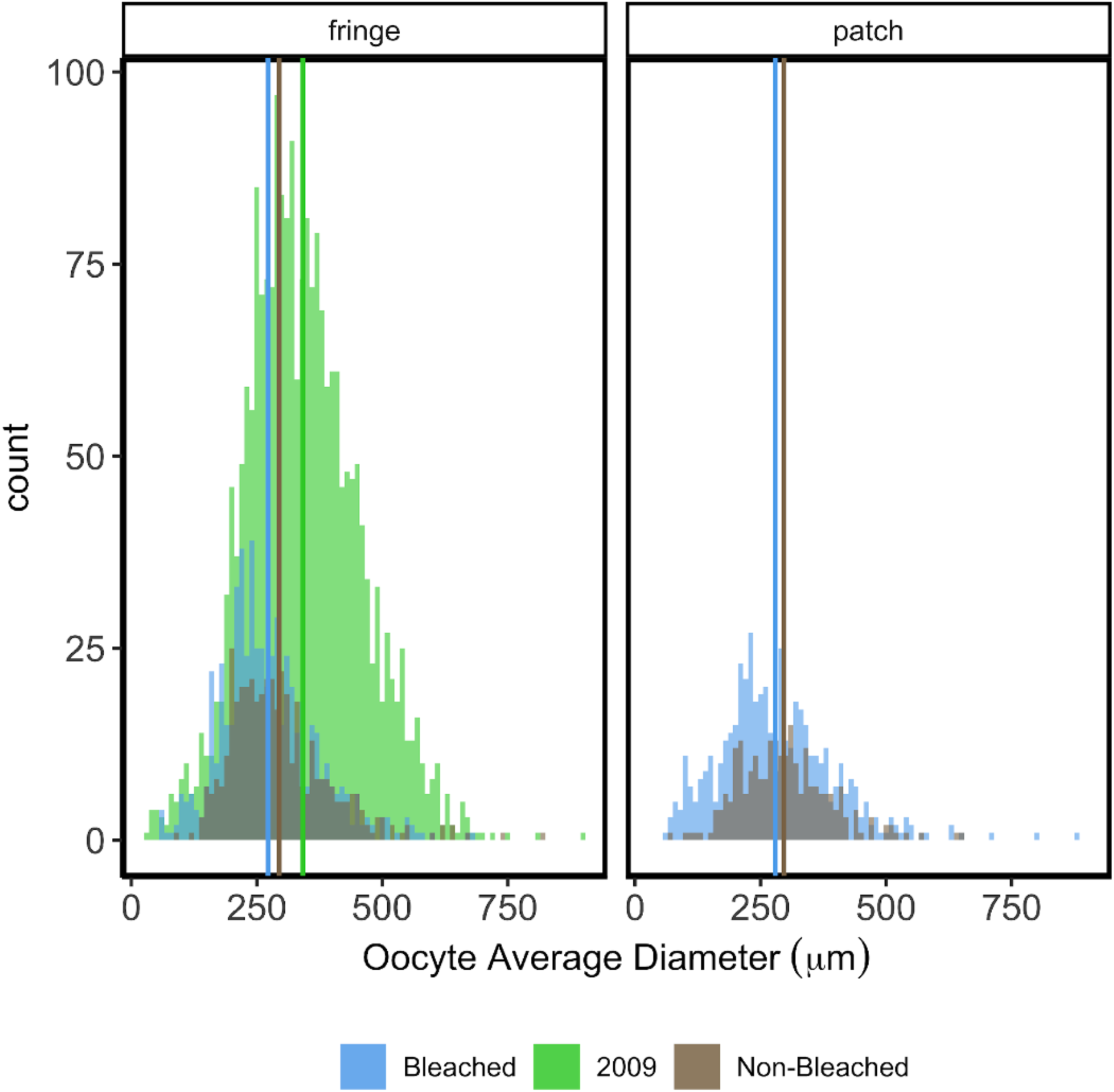
Oocyte size distribution in bleached and non-bleached colonies of *Montipora capitata*. Vertical lines represent mean values.

There was a significant difference in oocyte stages between reef type (p=0.0082) and bleaching history (p=7.0e-4) (Fig. 5). Oocytes were less developed (lower stage number) in colonies that bleached than in colonies that did not bleach. Fringing reefs had more developed oocytes (higher stage number) compared to patch reefs (Fig. 5).

**Figure 5.**
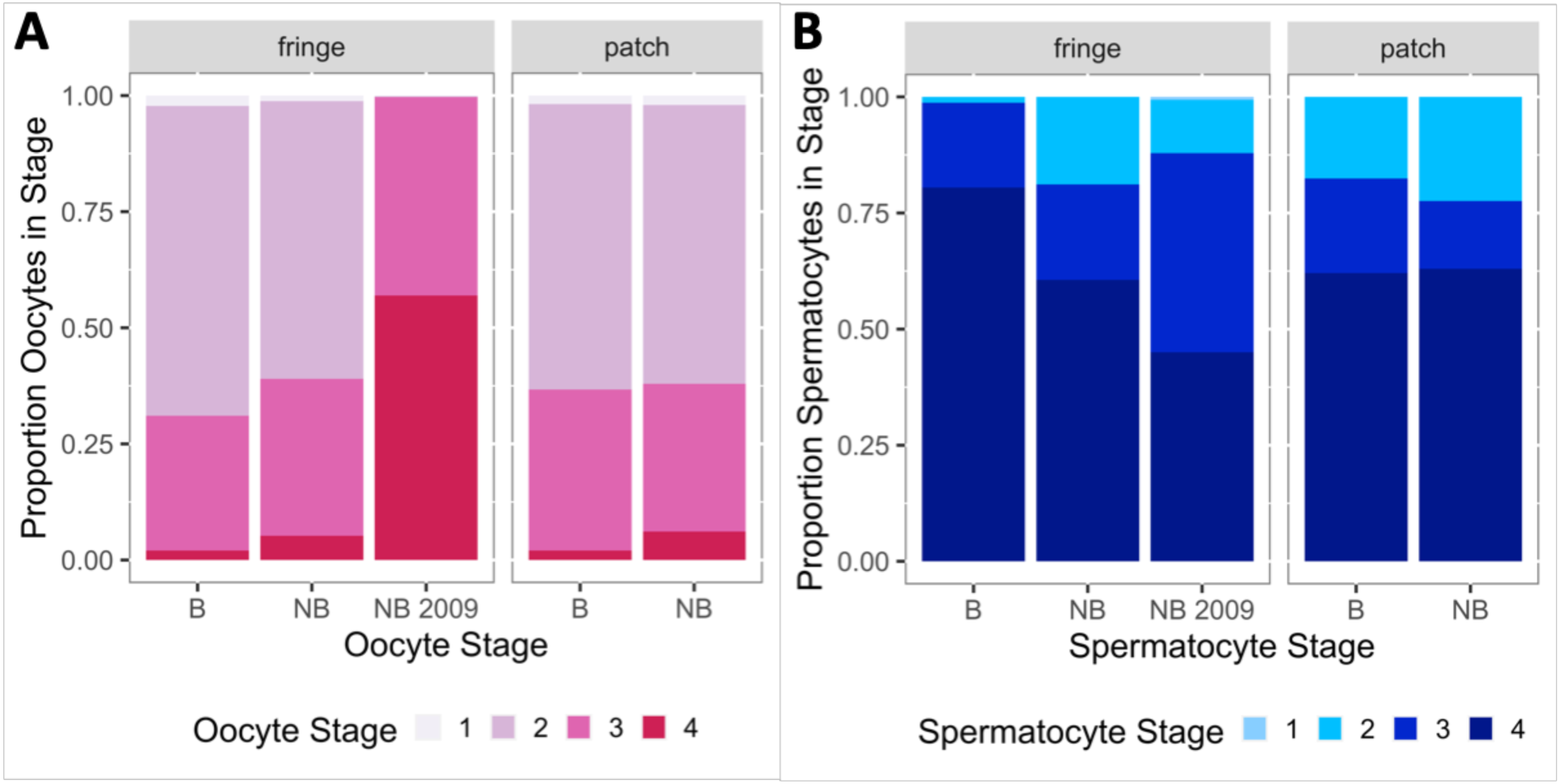
Oocyte and spermatocyst stages in bleached and non-bleached colonies of *Montipora capitata*

For the inter-year comparison, gamete stage of both oocytes and spermatocysts (comparing within fringing reef only) differed significantly by year (p= 2e-16 for both comparisons). The oocytes and spermatocysts measured in 2009 were at more advanced stages of development than those from non-bleached colonies in 2015. There was no significant difference in oocyte and spermatocyst stages based on previous bleaching or reef type (p>0.05, Fig. 5).

### Transcriptome assembly and annotation

Genome-guided transcriptome assembly resulted in 578,756 transcripts (N50 = 1,779bp) with median and mean lengths of 388bp and 861bp, respectively. There were 453,290 genes (N50 = 858bp) assembled with median and mean lengths of 328bp and 601bp, respectively (Supplementary Table 1). Transdecoder identified 142,601 complete, long open reading frames. Trinotate analysis resulted in 46,840 genes being annotated with Gene Ontology terms. The CD-hit to corset pipeline yielded a transcriptome with reduced redundancy and 339,648 transcript clusters. Almost 38% of these transcript clusters (n=128,570) were annotated with BLASTx against the UniProt trembl database (April 29, 2018).

### Differential Gene Expression Analysis

Weighted gene correlation network analysis was performed on transcripts detected in at least 10 out of 12 corals. Eleven WGCNA modules were strongly correlated with reef type (correlation coefficient > |0.7|) and four were strongly correlated with bleaching status (Supplemental Table 2). As with the differential expression analysis, there was a stronger signal of transcriptomic differences by reef type than by bleaching status in the WGCNA results. A complete list of enriched GO terms in the significantly correlated modules can be found in Supplementary Table 2. These included L-ascorbic acid biosynthetic process (yellow module, significant by reef type); sperm connecting piece (darkturquoise module, significant by reef type); cellular oxidant detoxification (blue3 module, significant by reef type); protein monoubiquitination (ivory2 module, significant by reef type); regulation of apoptotic process (hotpink module, significant by reef type); and calcium ion transmembrane transport (violetred4 module, significant by reef type).

#### Montipora reef type comparison

Ordination analyses (NMDS and discriminant analysis of principal components, DAPC) reveal a stronger, lasting impact of bleaching at the patch reef than at the fringing reef (Fig. 6). ANOSIM revealed a significant difference in transcriptomes by reef type (ANOSIM R = 0.4074; p = 0.012), but not by bleaching status (p>0.05). Transcripts with strong loadings on the DAPC LD1 axis, which separates bleached patch reef transcriptomes from others, included proteins involved in the cytoskeleton, skeletogenesis, extracellular matrix, and the immune response (Supp. Table 3). Transcripts with strong loadings on LD2, which separates non-bleached patch reef transcriptomes from others, included functions such as cholesterol homeostasis, immune response, bile acid transport, and cytoskeleton (Supp. Table 3).

**Figure 6.**
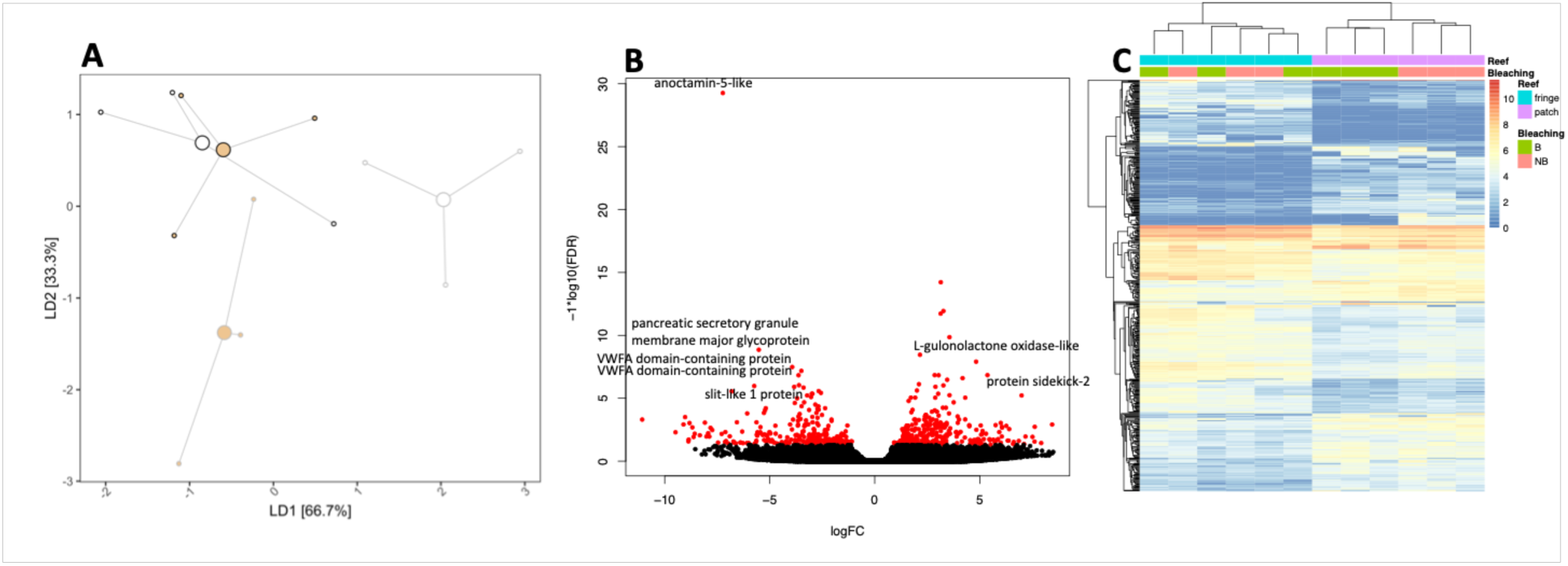
A. Discriminant analysis of principal components of bleached (white) and non-bleached (tan) M. capitata from patch reef (gray outline) and fringing reef (black outline). Transcripts loaded on LD1 represent 66.7% of the variation between groups; transcripts loaded on LD2 represent 33.3% of the variation between groups. B. Differentially expressed genes (DEGs) between reef types. The volcano plot shows the 563 DEGs in red; transcripts not differentially expressed are in black. Positive log fold change (LFC) transcripts were higher at the fringing reef and negative LFC transcripts were higher at the patch reef. C The heat map represents the DEG transcripts in each sample, with blue representing lower expression and red representing higher expression.

There were 563 differentially expressed genes in a comparison of corals (bleached and non-bleached) from the two reef types (Fig. 6), representing a wide range of enriched GO terms. Transcripts that were at higher abundance from the patch reef were enriched for GO terms including calcium transport, growth hormone signaling, organ development, and cell-matrix adhesion (Fig. 7). There were also categories of DEGs that were not included in the enrichment results but are of note. Several transcripts associated with calcium binding or calcium transport were elevated at the patch reef compared to the fringing reef. Multiple transcripts typically associated with embryonic development were also elevated, as were transcripts involved in the immune response and mucus production.

**Figure 7.**
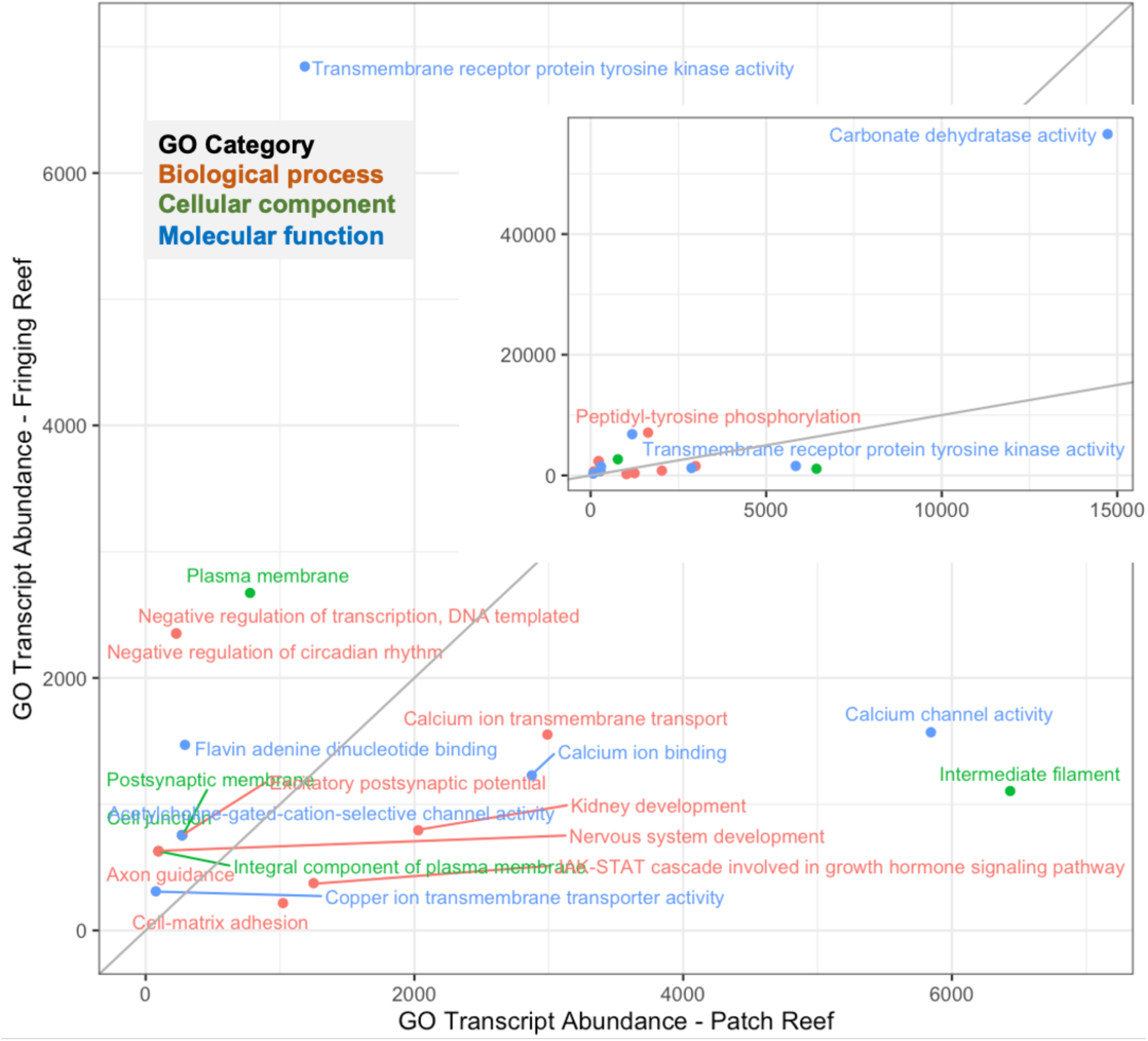
GO terms enriched in the significant DEGs between reef types. Transcript abundance for the patch reef for each GO term is plotted on the x-axis and abundance for the fringing reef is on the y-axis. Terms are colored by GO category. The gray line represents the 1:1 line such that GO terms lying above the line are at higher abundance in fringing reef transcriptomes and those below the line are at higher abundance from the patch reef. The inset graph is the entire dataset; the full plot is zoomed in to exclude the highly expressed carbonate dehydratase activity.

Transcripts that were at higher abundance in fringing reef corals were enriched for GO terms associated with nervous system, protein post-translational modification, and negative regulation of circadian rhythm (Fig. 7). These general differences between the sites suggest an underlying physiological difference between corals from different reef types. Categories of transcripts that were at higher abundance from the fringing reef, but not included in enrichment results, included apoptosis, extracellular matrix, protein metabolism and ubiquitination, multiple forms of carbonic anhydrase, transcription, circadian clock (Fig. 6 and 7), vitamin C biosynthesis (Supp. Fig. 1), and androgen metabolism.

#### Bleaching comparison

In a comparison of bleached and non-bleached corals, regardless of reef type, 32 transcripts were differentially expressed (Fig. 8). Transcripts that were at elevated levels in non-bleached corals were associated with energy transduction, sodium independent transport, sphingolipid metabolism, DNA replication, and immune response. Transcripts that were significantly higher in bleached corals were associated with glutathione catabolism, and DNA binding (Supplemental Table 3). Two transcripts (integrase catalytic domain-containing protein and reverse transcriptase domain-containing protein) may be microbial contamination because annotations suggest they are involved in insertion of viral and/or bacterial DNA into the host genome.

**Figure 8.**
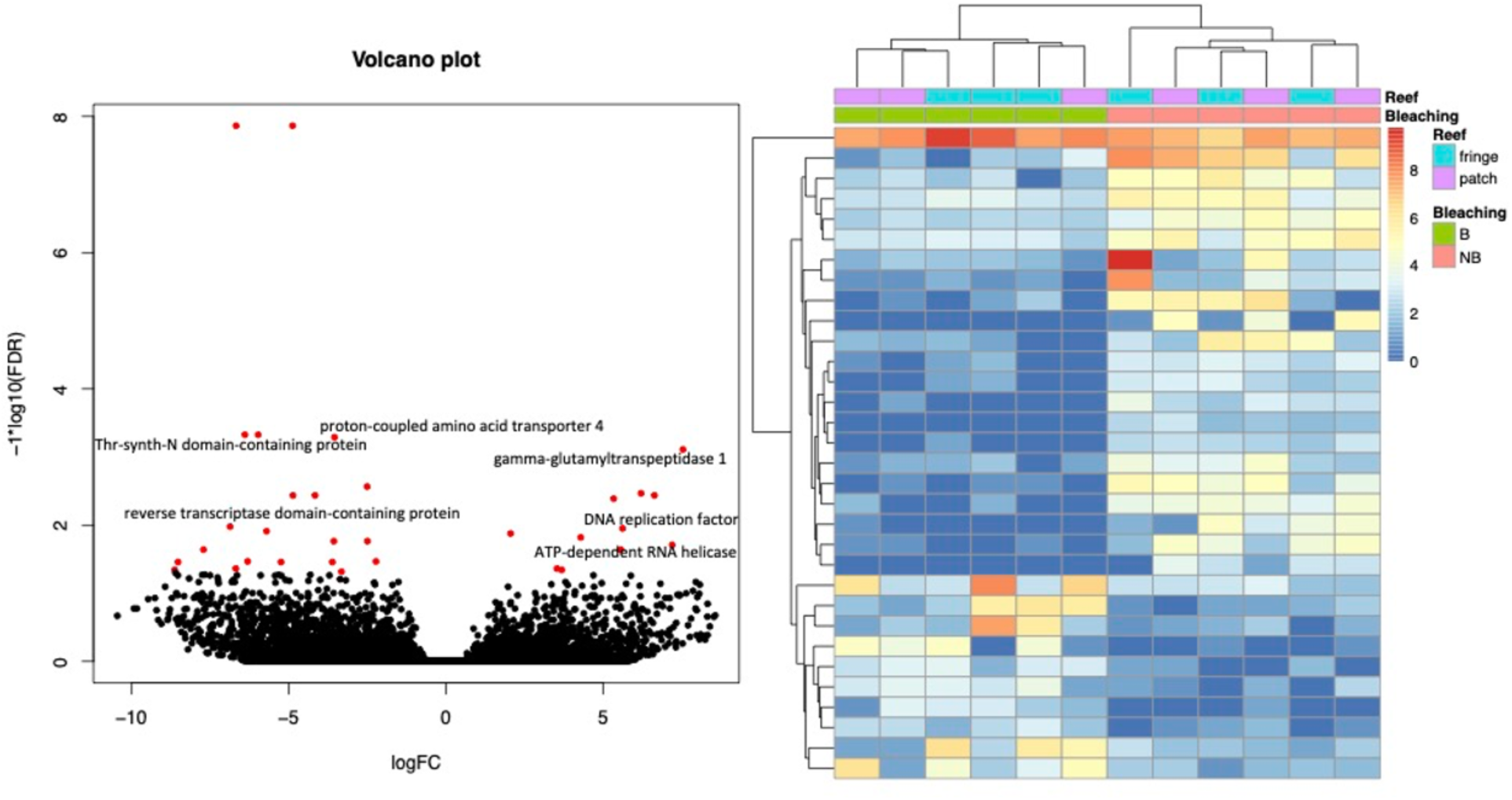
Differentially expressed genes between bleached and non-bleached corals. The volcano plot (left) shows the 32 DEGs in red; transcripts not differentially expressed are in black. Positive log fold change (LFC) transcripts were higher in bleached corals and negative LFC transcripts were higher in non-bleached. The heat map on the right represents the DEG transcripts in each sample, with blue representing lower expression and red representing higher expression.

#### Fringing Reef

In a comparison of bleached and non-bleached corals from the fringing reef only, 36 transcripts were differentially expressed (Supplemental Fig. 2, Supp. Table 3). Eight different transcripts representing the gene mitochondrial import receptor subunit TOM70 were differentially expressed between B and NB corals at site K4. These transcripts were elevated in non-bleached corals and are responsible for the enrichment of the two GO terms “Regulation of apoptotic process” and “Transferase activity, transferring glycosyl groups”. Other transcripts elevated in NB corals were associated with apoptosis and immune response. In bleached corals, the elevated transcripts included two annotated as pogo transposable element with KRAB domain (transcription) and CH2-type domain-containing protein.

#### Patch Reef

There were 57 DEGs in a comparison of bleached and non-bleached corals from the patch reef (Supp Table 3, Supplemental Fig. 3). Annotations of transcripts that were elevated in non-bleached corals revealed transcripts associated with extracellular matrix binding/axonal growth, binding of simple sugars, transfer of a sulfo group, amine as an acceptor, and DNA polymerase.

In bleached corals the following transcripts were elevated: tyrosine-protein phosphatase non-receptor type 12 (dephosphorylation), RNA-directed DNA polymerase from mobile element jockey (DNA replication), nose resistant to fluoxetine protein 6 (lipid transport, including yolk transport to oocytes), two transcripts of insoluble matrix shell protein, two transcripts representing RNA-directed DNA polymerase from transposon BS (DNA polymerase), 52 kDa repressor of the inhibitor of protein kinase (may inhibit cell growth), serine/threonine protein kinase pats1, GIY-YIG domain-containing protein (DNA repair nuclease).

## DISCUSSION

### Long-term impact of bleaching on coral reproduction

In this study we examined long-term impact on fitness and sublethal effects in *Montipora capitata* eight months after a natural bleaching event. We obtained colonies from fringing and patch reefs to test whether (1) gamete presence, fecundity, oocyte size and gametogenesis were affected by bleaching, (2) transcriptomic signatures differed between corals that bleached and did not bleach, and (3) environment influenced coral transcriptomic signatures and reproductive performance.

Our results show that reproduction in *M. capitata* is a resilient process but can be affected by thermal stress. We found that most colonies (100% in the fringing reefs and 81% in the patch reefs) were able to develop gametes despite bleaching, and that polyp fecundity did not change with bleaching history or between reef types. However, colonies that bleached had smaller oocytes and slower oocyte development than non-bleached colonies and this trend was consistent in both fringing and patch reefs. It is important to note that oocyte sizes of bleached and non-bleached colonies were smaller compared to oocytes from previous studies in years when no bleaching was reported (Cox, 2007; Padilla-Gamiño et al., 2014). This shows that the thermal stress event also affected oogenesis in colonies that did not show evident signs of bleaching.

The delay in gametogenesis in bleached colonies may be associated with the time it took corals to regain their symbiotic algae after thermal stress and recover physiologically. After the thermal stress (October 2014), bleached corals had lower chlorophyll levels and biomass (Wall et al., 2019, Ritson-Williams & Gates, 2020), possibly impacting the energy available to continue the development of the gametes. Oocyte development in *M. capitata* starts in the summer and oogenesis lasts 9-11 months (Padilla-Gamiño et al., 2014). Thermal stress has the potential to affect not only the adult colonies but also the oocytes that will be released the following summer. Rates of bleaching recovery in Hawaiian corals are on the order of months (Jokiel & Coles 1977, Bahr et al., 2015; Jokiel & Brown, 2004; Ritson-Williams & Gates, 2020; Rodrigues & Grottoli, 2007; Wall et al., 2019). During the year of our study (2014-2015), *M. capitata* colonies recovered chlorophyll levels and biomass throughout Kāneʻohe Bay by January, three months after the bleaching event (Wall et al. 2019).

It is interesting to note that lipid levels in *M. capitata* after the thermal stress (bleaching) were not affected (Wall et al., 2019). This is consistent with previous findings showing that bleached colonies of *M. capitata* can rely on heterotrophic feeding during recovery (Grottoli et al., 2006; Hughes et al., 2010; Rodrigues & Grottoli, 2007) and thus have the resources to prioritize gamete development despite loss of symbionts (Rodrigues & Padilla-Gamiño, 2022).

Spermatocyst development did not change between bleached and non-bleached colonies or reef types in 2015. Most of the colonies at fringing and patch reefs had spermatocysts at stage 4, which is the most mature stage and the development necessary for successful fertilization. Although not significant, bleached colonies in fringing reefs showed the highest proportion of colonies with spermatocysts at stage 4 (>75%). Spermatogenesis in *M. capitata* takes 4-5 months and occurs from April to August (Padilla-Gamiño et al., 2014). This process is less likely to be affected by thermal stress (compared to oogenesis) because it starts after bleached colonies have regained their symbionts. However, when we compare spermatogenesis between years with and without thermal stress (Padilla-Gamiño et al., 2014), we found that spermatocyst development may be faster in the year when corals bleached. In 2009, a year when no bleaching occurred, proportions of spermatocysts at stages 3 and 4 had very similar proportions (∼40%) compared to 2015, when most colonies had spermatocysts at stage 4 (Fig. 5b). Faster spermatocyst development in bleached years can be a problem especially if oocyte development is delayed, as our results suggest. If the spermatocysts develop faster than the oocytes, this can have important consequences for sperm quality and senescence, spawning synchronization, fertilization success, and offspring quality (Levitan & Petersen, 1995). In 2015, post-thermal stress, M. capitata sperm had a 44% motility reduction compared to the previous four years (normal range 70-90%), and larval survival and settlement were also compromised (Hagedorn et al., 2016). Furthermore, sperm motility remained less than 50% in 2017 and 2018 (years when bleaching also occurred) and had not fully recovered by 2019, with ∼63% motile sperm (Henley et al., 2021). This reduction in sperm motility in *M. capitata* was associated with mitochondrial damage but no molecular mechanisms were investigated (Henley et al., 2021). Future research is needed to better understand how thermal stress affects the plasticity and molecular mechanisms involved in the timing of gamete development, gamete quality and synchronization of their release.

### Transcriptomic profiles differ between bleached and non-bleached colonies eight months after a bleaching event

*M. capitata* expressed prolonged bleaching effects in coral colonies from both reef types, persisting eight months post-event. Our study revealed reef-specific transcriptomic signatures, indicating distinct long-term molecular impacts of bleaching. Previous work has shown that a lasting transcriptomic signature of coral bleaching can persist for up to 12 months after the event in other species (Thomas et al., 2019; Thomas & Palumbi, 2017); however, the specific transcriptomic response can vary by species (Thomas et al., 2019). Similarly, distinct metabolomic signatures have been observed in corals with different bleaching susceptibility four years post-bleaching (Roach et al., 2021).

Across coral species, the short-term transcriptomic bleaching response is usually defined by increased levels of transcripts associated with heat shock, oxidative stress, apoptosis, unfolded protein response and decreased abundances of transcripts associated with the extracellular matrix, calcification, and innate immunity (DeSalvo et al., 2008, 2010, Vidal-Dupiol et al., 2014; Ruiz-Jones & Palumbi, 2017; Thomas & Palumbi, 2017; Williams et al., 2021). In our study, some of these signals were still detected in *M. capitata* eight months after bleaching. Given the baseline differences in transcriptomes between reef types (see below), it is likely that *M. capitata*’s transcriptomic response to bleaching differs by location and may be impacted by local adaptation.

Despite little overlap in transcriptomic response to bleaching between reef types, bleached corals from both types of reefs had transcriptomic signatures suggesting negative impacts on sperm energy production and motility. Transcripts associated with mitochondrial creatine kinases (B,M,S and U-type) were suppressed in corals bleached across reef types. Creatine kinases are crucial in supplying energy for sperm motility (Dorsten et al., 1997), and are part of the *M. capitata* sperm transcriptome (Van Etten et al., 2020). The suppression of these genes likely led to reduced sperm motility in bleached corals (Henley et al., 2021), which could have implications for fertilization success. In the current study the mitochondrial import receptor subunit TOM70, which is involved in mitochondrial stress (Backes et al., 2021), was at lower abundance in bleached corals. Both TOM70 and mitochondrial creatine kinases may be important biomarkers of bleaching stress, and the link between bleaching, the expression of genes, and sperm function should be further explored.

Despite little overlap in reef-specific responses to bleaching stress, two transcripts were differentially abundant between B and NB corals at both reefs: an interferon-induced very large GTPase and a transcript with no annotation. Interferon-induced very large GTPases are immune response proteins. A lingering signal of altered expression of immune response transcripts is a common theme across the studies that have followed long-term bleaching responses (Pinzón et al., 2015; Thomas & Palumbi, 2017). The universality of the interferon-induced GTPase as a molecular signal of long-term bleaching stress, and its impact on coral immune function, is an additional avenue of further research.

### Reef-specific responses to bleaching

Transcripts involved in diverse physiological processes sustained altered expression levels in corals that bleached from the patch reef, including those involved in DNA replication and repair, lipid transport, protein dephosphorylation, and shell matrix. Transcripts that remained suppressed were involved in the extracellular matrix (ECM), binding of simple sugars, amine sulfotransferase, and DNA polymerase. ECM transcripts have been found to be impacted in the short term by thermal stress and bleaching (DeSalvo et al., 2008), and it is hypothesized that their downregulation could impact calcification. Additionally, transcripts with strong loadings in the DAPC analysis associated with bleaching status at the patch reef included those annotated as carbonic anhydrase, mannose-binding protein C, and lymphocyte antigen 6D. Carbonic anhydrase may be associated with coral skeletogenesis (Bertucci et al., 2013) and is further evidence that calcification and somatic growth may remain impacted in corals at the patch reef. The two immune-related transcripts - mannose-binding protein C and lymphocyte antigen 6D - associated with bleaching at the patch reef may be biomarkers of lasting bleaching impacts on these coral’s innate immune system. Mannose-binding lectin appears to be a biomarker of the thermal stress response in corals and may be necessary in maintaining symbiosis (Vidal-Dupiol et al., 2009). Its transcript was up-regulated in thermally preconditioned corals in response to a thermal challenge (Bellantuono et al., 2012) and was down-regulated during thermal stress and bleaching in *P. damicornis* (Vidal-Dupiol et al., 2009). These pieces of evidence suggest that this transcript, and its protein, may be protective against bleaching.

At the fringing reef, transcripts that maintained altered expression levels eight months post-bleaching represented a distinct set of processes compared to the patch reef. Transcripts that were elevated in bleached colonies included those involved in transcription. Transcripts that were still suppressed included the processes of apoptosis, sphingolipid metabolism, serine hydrolases, and the immune response. The suppression of transcripts associated with apoptosis and the immune response is a typical short-term coral response to thermally induced bleaching (e.g., Vidal-Dupiol et al., 2014), and yet, in *M. capitata*, it remains a biomarker of bleaching at the fringing reef months after the stress-inducing event. Similarly, genes regulating apoptosis and immunity were also down-regulated up to a year after bleaching in *O. faveolata* (Pinzón et al., 2015); however, in *Acropora hyacinthus*, functionally similar transcripts were up-regulated and remained so for months after bleaching (Thomas & Palumbi, 2017). The cellular and physiological stress induced by a bleaching event seems to have lasting impacts on the coral immune response, but the specific molecular markers (and perhaps physiological details) of those impacts vary by species and perhaps also by location. In the short term, this bleaching-induced weakening of the immune response has proven to make *P. damicornis* more susceptible to infection in a lab setting (Vidal-Dupiol et al., 2014); it remains to be seen if long-term impacts on the immune response affect susceptibility to infection in different species in the field.

### Corals had site-specific transcriptomic profiles

There are innate molecular differences between *M. capitata* found at the two reef types, patch and fringing. These site-specific differences were a driving factor across statistical analyses suggesting that *M. capitata* is acclimatized and/or adapted to its local environment. Reef type-specific differences in the transcriptome are also reflected in the lingering molecular signatures of bleaching, which had very little overlap in specific transcripts or in overall transcript function between the two reef types. These reef type-specific transcriptomic profiles may give insight into local environmental drivers that shape coral physiology and environmental response.

Differentially abundant transcripts between reef types represent a wide range of functions, highlighting the complex nature of local acclimatization/adaptation and environmental response. A strong, reef type-specific transcriptomic signature has previously been detected in *M. capitata* (Helmkampf et al., 2019) and in *Montastraea cavernosa* (Studivan & Voss, 2020). Many of these transcripts that drive the transcriptomic differences between colonies from different reefs may also be involved in the different schedules of bleaching recovery at the two reefs. Immune response transcripts had a higher abundance in patch reef corals. The expression levels of immune response transcripts are typically altered in response to various biotic and abiotic stressors, such as growth anomaly disease (Frazier et al., 2017), warming (Williams et al., 2021), bleaching (Thomas & Palumbi, 2017), and simulated climate change (Kaniewska et al., 2015), suggesting that patch reef corals may experience a higher level of environmental stress than fringing reef corals. Calcium transport transcripts were also revealed as an important driver in transcriptomic differences between reef types. It may be that somatic/skeletal growth is prioritized in the patch reef corals, whereas gametogenesis is prioritized in the fringing reef corals. The link between higher environmental stress and increased calcification/somatic growth, along with other physiological processes that differ by reef type, may reveal key elements to site-specific stress response and survival in *M. capitata*.

Additional evidence of an elevated physiological stress response in patch reef corals, and higher rates of somatic growth, comes from the elevated transcripts frizzled (Wnt receptor) and notch. These transcripts are environmentally sensitive in corals and have also been found to be differentially abundant in response to growth anomaly disease (Frazier et al., 2017), low pH (González-Pech et al., 2017) and simulated climate change (Kaniewska et al., 2015). The Wnt pathway genes, including frizzled receptors, are highly conserved across metazoa. This pathway is essential in embryonic patterning and gastrulation across taxa, and this essential role has been confirmed in multiple taxa of cnidaria (Lapébie et al., 2014; Guder et al., 2006). However, the Wnt pathway also plays an important role in adult tissues in cnidaria, with genes in the pathway both constitutively expressed in localized tissues (Sanders & Cartwright, 2015) and up-regulated during tissue turnover and repair (Guder et al., 2006; Loh et al., 2016). The elevated expression of these transcripts in the patch reef corals may suggest increased tissue growth due to damage or a higher investment in somatic growth.

Coral colonies from fringing reefs exhibit quicker reproductive and transcriptomic recovery compared to those from patch reefs. This accelerated recovery may be attributed to the heightened environmental variability at fringing reef sites, enhancing the corals’ acclimatization capacity. In fringing reefs, we found a higher likelihood of gametes being present and oocytes being more developed than in patch reefs. Transcriptomic signatures indicate a closer resemblance between bleached and non-bleached colonies at fringing reefs, suggesting a faster recovery process compared to patch reefs. Additionally, fringing reef corals show transcriptomic signals associated with elevated skeletogenesis compared to their counterparts in patch reefs.

Many of the processes represented by higher transcript levels in fringing reef corals have proven to be environmentally sensitive in other coral transcriptomic studies, including carbonic anhydrase and vitamin C metabolism. Carbonic anhydrases perform two main roles in corals: 1) acquisition of CO_2_ for dinoflagellate symbiont photosynthesis (Tansik et al., 2015; Weis et al., 1989) and 2) skeleton calcification (Moya et al., 2008). It is difficult to ascribe a specific role to the carbonic anhydrases detected in this set of DEGs at the fringing reef; however, both processes represent positive coral metabolism and growth. The increased abundance of vitamin C synthesis transcripts lends additional evidence for increased skeletal growth in fringing reef corals. Vitamin C transport transcripts have previously been associated with coral calcification (Caspasso et al. 2022). L-gulonolactone oxidase in the vitamin C (ascorbic acid) biosynthesis pathway was at increased abundance in the corals from the fringing reef, lending additional evidence to the hypothesis of elevated calcification in the fringing reef.

Transcripts in apoptotic pathways were relatively elevated in fringing corals and have been found to be elevated across coral species exposed to various environmental drivers. The apoptopic pathway is a biomarker of coral environmental response; as coral proteins and cells sustain damage, apoptosis may be up-regulated to clear damaged proteins and cells. An increase in transcripts linked to apoptotic pathways is frequently detected at elevated levels post-environmental stress in corals (Cleves et al., 2020; Davies et al., 2016; Kaniewska et al., 2015; Pinzón et al., 2015; Thomas & Palumbi, 2017; Zhang et al., 2022). Similar transcriptomic signatures indicating protein damage or ubiquitination have been consistently observed in previous studies examining coral environmental responses (e.g., (Pinzón et al., 2015; Ruiz-Jones & Palumbi, 2017). These are likely signals of endoplasmic reticulum (ER) stress and reactive oxygen species damage as coral cells respond to environmental stresses (e.g. Petrou et al., 2021; Roach et al., 2021). Some corals change the abundance of ER stress transcripts during cyclical, sub-bleaching environmental variability (Ruiz-Jones & Palumbi, 2017), and the elevated levels of these transcripts could be an artifact of sampling day/time at the fringing reef.

### Conclusions/Broader Implications

The combined approach of assessing bleaching response and recovery using physiological (gametogenic) and transcriptomic responses revealed important links between cellular priorities and organism-scale reproductive processes. Even though gametogenesis progressed in all corals, oocyte development was delayed in bleached colonies. This can have important implications for synchrony in gamete release, fertilization, offspring development and performance.

We used transcriptomics to examine the mechanisms of physiological resilience and long-term impacts of bleaching. Our data show that corals with different bleaching susceptibility have contrasting transcriptomes eight months following a natural bleaching event. Suppression of sperm motility, calcification and immune-related transcripts were associated with bleaching susceptibility. We also found distinct transcriptomic profiles between fringing and patch reefs, indicating acclimatization and/or local adaptation in *M. capitata*. It is noteworthy that, following an 8-month period of recovery after thermal stress, the transcriptomic profiles of bleached and non-bleached colonies within the fringing reefs exhibited a greater degree of similarity compared to those in the patch reef, suggesting a faster physiological recovery rate among colonies from the fringing reef. Since colonies in the fringing reef are exposed to higher environmental variability, they may have evolved or have acquired plasticity to recover from stress more rapidly. This faster recovery extends to gametogenesis: colonies from the fringing reef were more likely to have gametes, which were more developed than those from the patch reef.

To protect coral reefs and better understand how coral populations will be affected by future thermal stress, we need to know not only which colonies survive but also how their physiological and reproductive processes are affected in the short and long-term. It is essential to understand the dynamics of recovery as consecutive bleaching events are becoming more frequent, potentially limiting recovery time. Our study examines the variability in long-term recovery and provides molecular and reproductive data that can aid in conservation and management efforts. These insights can help to detect biomarkers of bleaching susceptibility and resilience, project the impact of future bleaching events on reproductive output, assess which traits are more vulnerable, and identify what sites are more likely to be compromised.

## ACKNOWLEDGEMENTS

Special thanks to CSUDH students Ashley Trujillo, Janet Mejia, Marshay Galloway, Vanessa Gomez Mendez and Azia Mitchell. Thanks to the Gates Lab and HIMB staff for their support in the field. Thanks to Holly Clark, Ann Tarrant and Amy Maas for guidance on transcriptomic analysis. This research was conducted under Hawai’i Department of Land and Natural Resources scientific permit 2017-13 to JPG.

The data that support the findings of this study are openly available in https://github.com/Nunn-Lab/Publication-Mcapitata-bleaching-transcriptomics

Data available:

Egg and sperm development (gamete stages), egg size, polyp fecundity (eggs per polyp), transcript counts.

