## Supplemental figures and text for "Coral long-term recovery after bleaching: implications for sexual reproduction and physiology"

**Supplemental Figure 1**

**Genes in the ascorbate (vitamin C) metabolism pathway detected in the *M. capitata* transcriptome. L-gulonolactone oxidase (L-gulonolactone ox.) was differentially expressed at higher abundance from the fringing reef and is colored red. All other identified transcripts, but not differentially abundant, are colored in yellow.**

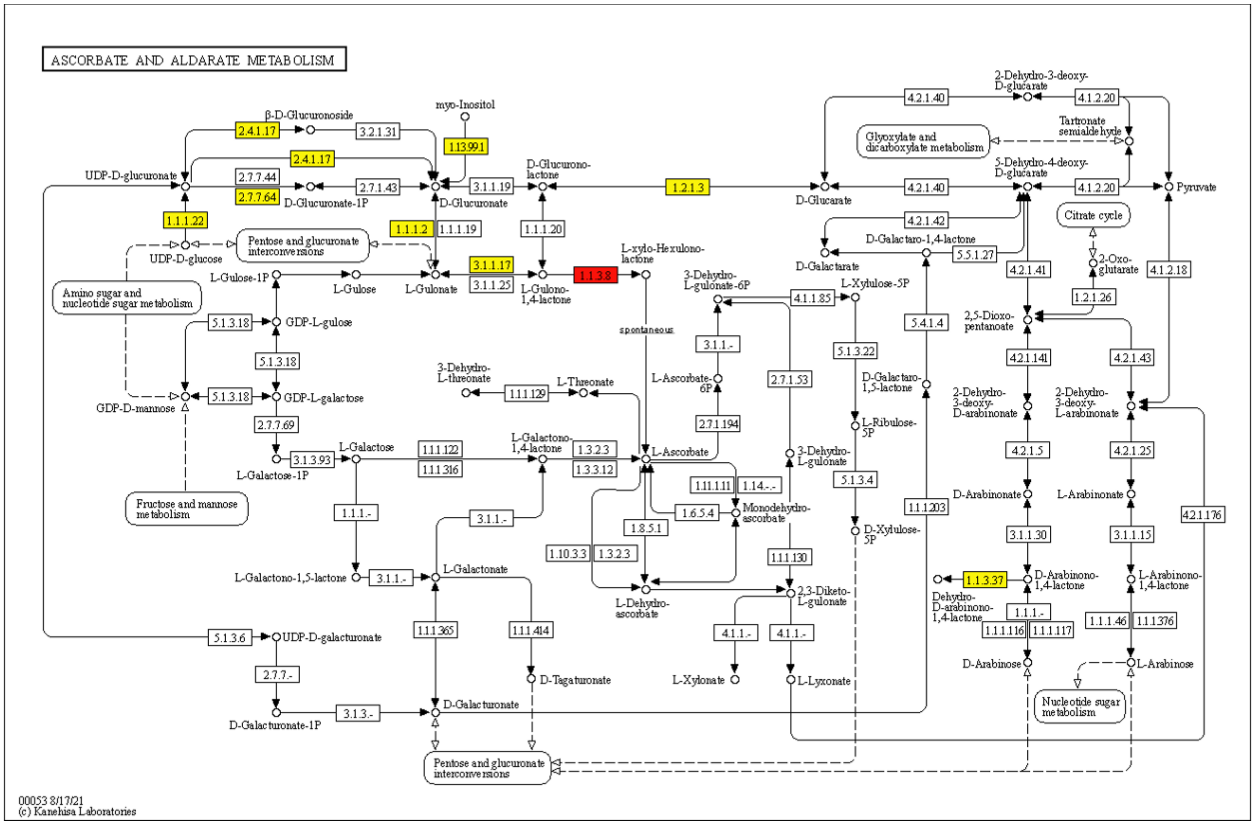


**Supplemental Figure 2**

**Differentially expressed genes between bleached and non-bleached corals from the fringing reef. The volcano plot (left) shows the 36 DEGs in red; transcripts not differentially expressed are in black. Positive log fold change (LFC) transcripts were higher in bleached corals, and negative LFC transcripts were higher in non-bleached. The heat map on the right represents the DEG transcripts in each sample, with blue representing lower expression and red representing higher expression.**

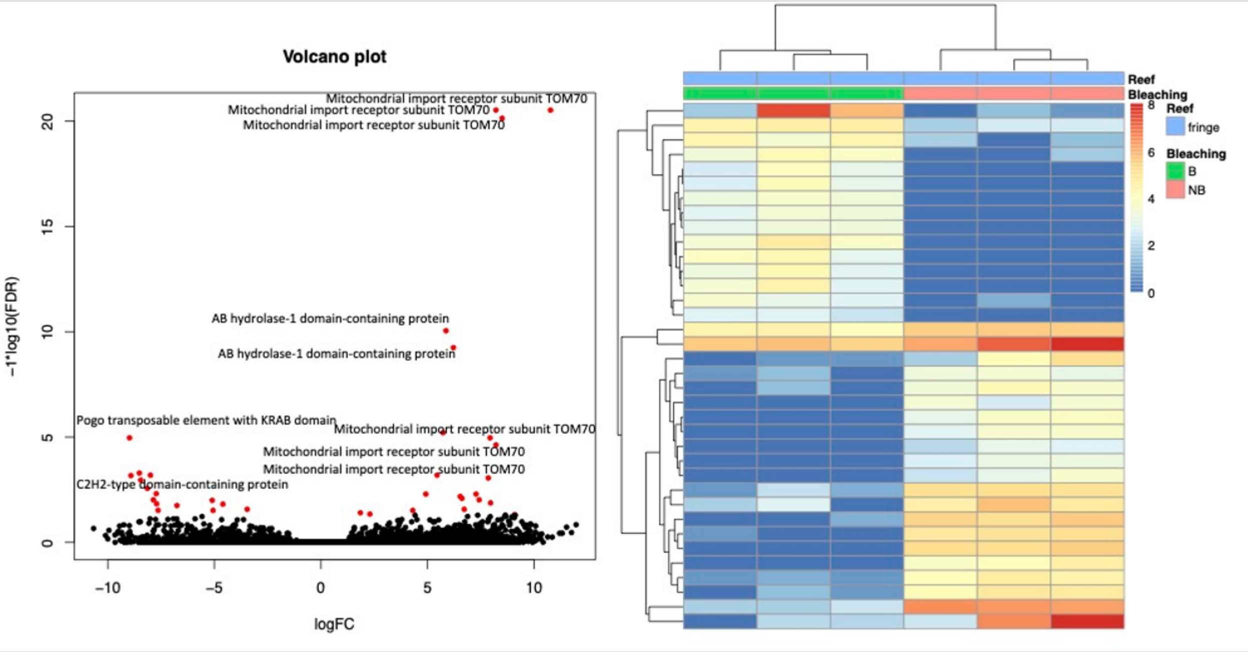

**Supplemental Figure 3**

**Differentially expressed genes between bleached and non-bleached corals from the patch reef. The volcano plot (left) shows the 36 DEGs in red; transcripts not differentially expressed are in black. Positive log fold change (LFC) transcripts were higher in bleached corals, and negative LFC transcripts were higher in non-bleached. The heat map on the right represents the DEG transcripts in each sample, with blue representing lower expression and red representing higher expression.**

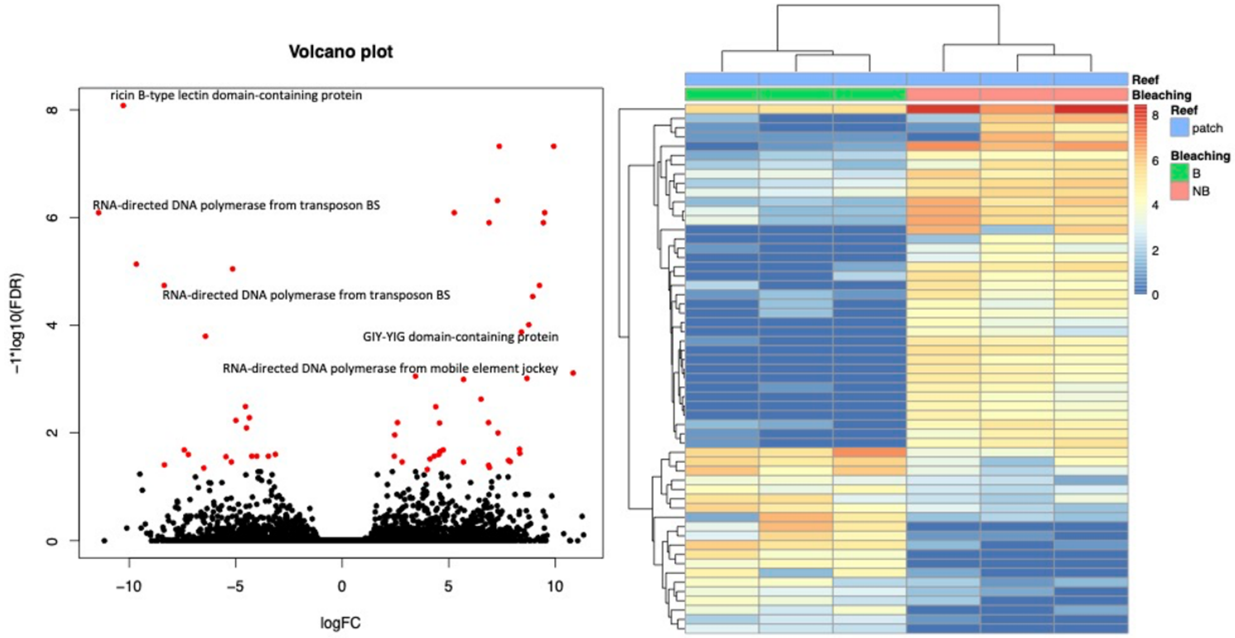

**Supplemental Methods and Analysis**

All FastQ files were quality trimmed, and adaptors removed, with Trim Galore! (v0.4.4_dev; https://www.bioinformatics.babraham.ac.uk/projects/trim_galore/) using Cutadapt (v1.16; Martin, 20111) and evaluated with FastQC (v0.11.7; http://www.bioinformatics.babraham.ac.uk/projects/fastqc/). A visual summary of the FastQC data was performed with MultiQC (v1.4; Ewels et al., 2016). A genome-guided transcriptome was *de novo* assembled with Trinity (v2.6.6; Grabherr et al., 2011, Haas et al., 2013) using trimmed reads. Transcriptome "completeness" was assessed with BUSCO (v3.1.0; Simão et al., 2015; Waterhouse et al., 2017) using the transcriptome option, metazoa_odb9 database, and AUGUSTUS (v3.3.2; Stanke et al., 2008; Stanke et al., 2006a; Stanke et al., 2006b; Stanke and Waack, 2003; Stanke 2003) with species set to "fly". TransDecoder (v5.5.0; Grabherr MG et al., 2011; Haas BJ et al., 2011) was used to identify putative open reading frames (ORFs), using BLASTp (v2.8.1+; Altschul et al., 1990, Camacho et al., 2008), and HMMER (v.3.2.1; http://hmmer.org/). Trinotate (v3.3.1; Bryant DM et al., 2017) was used to assign functional annotations to genes, using BLASTx (v2.8.1+; Altschul et al., 1990; Camacho et al., 2008), RNAMMER (v1.2; http://www.cbs.dtu.dk/services/RNAmmer/), SignalP (v4.1; Petersen et al., 2011), tmhmm (v2.0c; Sonnhammer ELL et al., 1998) and the longest ORFs identified by Transdecoder.

Redundancy in the transcriptome was reduced by applying the CD-hit-to-corset pipeline. CD-hit-EST (v. 4.8.1) was run on the Trinity assembly (453,290 transcripts) with a 0.98 similarity cut-off. The output from CD-hit-EST was used to build an alignment database in bowtie2 (v.2.4.4). Sequence alignment was performed with bowtie2 and resulting .bam files were run through corset (v. 1.0.9) to further reduce sequence redundancy into transcript clusters. Corset groups were set according to bleaching status and site (4 groups). Transcripts were reduced to 339,648 transcript clusters for downstream analysis.

**STATISTICAL ANALYSES**

**Reproductive status and bleaching**

Individual sites of colony collection were combined to look at overall reef type impacts, rather than individual site impacts. In all models, bleaching condition and reef type were fixed effects and colony was set as a random effect.

Gamete presence/absence was explored using a generalized linear mixed-effects model with a binomial response. Fecundity (oocytes per polyp), number of oocytes per stage and number of spermatocysts per stage were explored using a generalized linear mixed-effects model with a Poisson regression. Oocyte Feret diameter was explored with a linear mixed-effects model.

**Transcriptomics**

Only transcripts detected in at least 10 samples were included in the analysis. The counts data were normalized using median localization normalization before WGCNA analysis. We investigated the influence of reef type (fringing or patch) and bleaching status on the expression trends of groups of genes.

As part of the compGO pipeline, all *M. capitata* transcripts were searched against the UniProt Trembl database using DIAMOND. These annotations were compared with annotations resulting from BLASTx against UniProt/SwissProt and a consensus annotation was selected for all DEGs. To generate pathway annotations, DEGs were translated using TransDecoder and searched against the Kyoto Encyclopedia of Genes and Genomes using BlastKOALA with taxonomy of genome set to Eukaryotes and searched against the family_eukaryotes KEGG database. KEGG Mapper Color was used to illustrate specific pathways.
